# Stable primary brain cell cultures from zebrafish reveal hyperproliferation of non-neuronal cells from *scn1lab* mutants

**DOI:** 10.1101/2024.10.01.616210

**Authors:** G. Lorenzo Odierna, Sarah Stednitz, April Pruitt, Joshua Arnold, Ellen J. Hoffman, Ethan K. Scott

**Affiliations:** Queensland Brain Institute, The University of Queensland, St Lucia, Queensland, Australia; Child Study Center, Program on Neurogenetics, Yale School of Medicine, Yale University, New Haven, CT, United States; Department of Neuroscience, Yale School of Medicine, Yale University, New Haven, CT, United States; Department of Anatomy and Physiology, The University of Melbourne, Australia

**Author notes:** Corresponding author at current address: Menzies Institute for Medical Research, The University of Tasmania, Hobart, Australia.

**Keywords:** Zebrafish, Primary Culture, Neuron, Calcium Imaging, Sodium Channel, Hyperproliferation, *Scn1*

## Abstract

Zebrafish are a popular model system for studying the genetic and neural underpinnings of perception and behavior, both in wild-type animals and in the context of disease modelling. Cultured primary neurons provide a key complementary tool for such studies, but existing protocols for culturing embryonic zebrafish primary neurons are limited by short cell survival and low neuronal purity. In this study, we set out to establish a protocol to produce long lived, pure neuronal cultures from zebrafish that could be used to study the mechanistic contributions of genes to neuronal networks. We then used these primary cultures to characterize cell proliferation and differentiation in primary neurons derived from *scn1lab* mutant embryos, which lack a sodium channel relevant to Dravet syndrome and autism. Using our optimized protocol, we generated cultures that proliferate, diversify, and form stable networks of neurons surviving for months. These stable cultures allowed us to perform genetic experiments, in this case revealing dramatic differences in the cellular composition of cultures derived from *scn1lab* mutant embryos versus their wild type siblings. Specifically, we find that loss of *scn1lab* promotes hyperproliferation of non-neuronal cells in mixed cultures of brain cells. In pure neuronal cultures, we find alterations in neurotransmitter subtypes consistent with known effects of *scn1lab* loss of function. Validating the utility of this approach, we then identify a corresponding hyperproliferation phenotype in live *scn1lab* mutant embryos, shedding light on potential mechanisms that may be relevant for Dravet syndrome.

**Significance statement:** Most existing embryonic zebrafish primary neuron culture protocols describe growing mixed cell types for restricted durations. Here, we report generation of zebrafish mixed type or pure neuronal cultures that are viable for over 100 days. We apply these cultures to gain new insight into *scn1lab*, a zebrafish orthologue of the Dravet Syndrome-associated sodium channel gene *SCN1A*. We report that loss of *scn1lab* results in hyperproliferation of non-neuronal cells, revealing an underappreciated mechanism by which mutations in *SCN1A* impact the structure and function of neuronal networks. Our *in vitro* cultures thus faithfully recapitulate *in vivo* neurobiology and provide a powerful platform to interrogate brain function in health and disease.

## Introduction

Primary brain cell culture provides a valuable window through which insights into cellular and molecular mechanisms of brain development and function can be gained. This is the especially the case with small animal models, where cultures can readily be grown from organisms harboring targeted genetic mutations to complement *in vivo* observations. Despite this appeal, the culturing of zebrafish primary neurons has seen limited improvement since its initial application over two decades ago (Andersen, 2001). Cultures have been successfully grown from larval and adult brains, but a standardized protocol for the consistent generation of pure, long lived primary neuronal cultures from zebrafish does not yet exist. A common approach for collecting zebrafish brain cells involves enzymatically dissociating whole embryos or larvae (Acosta et al., 2018, Fan et al., 2007, Russo et al., 2018, Sakowski et al., 2012, Sassen et al., 2017). Although this approach’s technical simplicity makes it appealing, neurons in the resulting cultures rarely survive for experimental purposes beyond a few days and are not suitable for the study of neuronal physiology due to interference from other primary cell types. Some studies have addressed these issues by isolating brain tissue by dissection prior to dissociation (Andersen, 2001, Kroehne et al., 2017, Lopez-Ramirez et al., 2016, Meade et al., 2019, Patel et al., 2019, Tapanes-Castillo et al., 2014, Chen et al., 2013b, Fassier et al., 2010), using cell sorting technologies (Kinikoglu et al., 2014, Welzel et al., 2015), or enriching neuronal populations using culture media that preferentially promote neuronal survival and outgrowth (Taylor and Houart, 2024). Although these purifying steps increase the longevity of cultures in some cases, whether neurons in these cultures form mature functioning networks comparable to those seen using mammalian cell lines remains uncertain. It is clear, therefore, that although the zebrafish community has access to protocols for culturing neurons, there remains room for improvement. Having a reliable method to produce long lived, pure neuronal networks *in vitro* from zebrafish brains would serve a useful complementary method alongside more traditional strengths of the model system such as *in vivo* neural development and neural circuit analysis (Vanwalleghem et al., 2018, Simmich et al., 2012).

In this paper, we outline a new protocol for culturing embryonic zebrafish brain cells. We have optimized several factors in this protocol, providing meaningful improvements in the quality and longevity of cultured zebrafish neurons, and producing cultures that closely replicate important elements of embryonic brains *in vivo*. We present an analytical framework for measuring several characteristics of these cultures, including the prevalence of neurons, and neurotransmitter subtypes. To test the utility of this new approach, we cultured brain cells from fish lacking the voltage-gated sodium channel subunit encoded by *scn1lab*. We found an overrepresentation of non-neuronal cells in these cultures, and using this targeted result, confirmed that the same imbalance exists in larva brains *in vivo*. Overall, this protocol raises embryonic zebrafish primary neuronal culture to a similar standard seen in other systems and provides a new approach for the *in vitro* study of neurodevelopmental processes.

## Methods

### Zebrafish handling

All work was carried out in accordance with animal ethics approval SBS/341/19 from the University of Queensland Animal Welfare Unit. Adult zebrafish (*Danio rerio*) were raised, maintained and fed under standard conditions (Westerfield, 2000). All animals used in this study were on a TL *nacre* background. The *scn1lab* Δ44 mutation has been previously described (Kroll et al., 2021). TL *nacre* fish harboring *HuC:H2B-GCaMP6s* have also been described previously and used to quantify neuronal activity in zebrafish harboring mutations associated with neurodevelopmental disorders (Chen et al., 2013a, Constantin et al., 2020, Marquez-Legorreta et al., 2022, Freeman et al., 2014). Generation of the *gad1b:GAL4* BAC transgenic line is described in Förster et al. (2017).

After collecting and washing embryos, we raised them at 33°C to accelerate their development such that they were ready for experiments at 1 day post-fertilization (dpf). The amount of time the embryos spent at 33°C varied depending on when they were fertilized and when we aimed to dissect them the next day. We found that using the formula developed by Kimmel et al. (1995), H_T_ = h/0.055T-0.57 (where H_T_ = hours of development at temperature T, and h = hours of development to reach the stage at 28.5°C), reliably allowed us to reach a standard developmental time of 30-36 hours post fertilization (hpf) with variable time at 33°C. This was verified according to the phenotypic characteristics consistent with the Prim-15 to Prim-25 window such as ∼90° head-to-trunk angle, increased eye pigmentation and yolk extension length ≥ yolk ball diameter, but not by a factor of 2 (Kimmel et al., 1995). For experiments with *scn1lab* mutant animals, manually dechorionated embryos were genotyped on the day of experimentation by clipping a small portion of their tails with fine forceps. We allowed them to continue developing in cell culture media (described in *Plating, media, and incubation*) during the DNA extraction and PCR process. Embryos that were injured by the tail clipping procedure underwent obvious discoloration and were discarded. As inclusion criteria, we only used apparently healthy organisms that retained the touch responsivity and spontaneous coiling expected at their developmental stage.

### Generation of cultures

Embryos were manually dechorionated before dissections. All embryos and larvae were cold-immobilized on an ice slurry for at least 10 minutes before dissection. Dissections were carried out in sylgard-bottomed dishes within a drop of calcium- and magnesium-free Hank’s Buffered Salt Solution (HBSS). Brains from animals up to 6 dpf were manually dissected using fine forceps with the aims of reducing damage to brain tissue before dissociation and minimizing the number of non-neuronal cells included in the suspension. We found that we could reliably remove retinal tissue and a significant fraction of dermal tissue surrounding the brain for both embryos and larvae, but for embryos, this often came at the cost of losing the olfactory bulbs and parts of the developing telencephalon. Despite removal of dermal tissue, we cannot eliminate that possibility that cells derived from this tissue, such as fibroblasts, were included in the plated cell suspension. We also could not confidently isolate the hindbrain from the developing spinal cord for embryos, and as such, dissections of single embryonic brains included a variable number of rhombomeres. Single brains were transferred in a HBSS droplet on forceps to polypropylene PCR tubes containing 200 µl of seeding media (see *Plating, media, and incubation*) on ice while dissections continued. Once at least 10 brains had been dissected, we pooled the brains and washed them three times by removing solution (leaving at least 100 µl behind) and adding HBSS up to 600 µl each time. The solution was then removed from the brains (as much as possible without disturbing the brains) and gently replaced with 100 µl of room temperature Accumax (Stem Cell Technologies, Vancouver, Canada). The brains were left to incubate in Accumax at 28°C for 20 minutes, after which the Accumax was removed and 600 µl of seeding media was gently added. The sample was spun at 100 x relative centrifugal force (RCF) for 1 minute to form a pseudo pellet. Supernatant was removed without disturbing the pseudo pellet, and 200 µl of seeding media was added. Brains were then triturated rapidly 100 times using 200 µl plastic pipette tips set to 100 µl volume. A 10 µl sample of the suspension was taken for cell counting by mixing with 10 µl of trypan blue before using the cells for plating. Cells were counted via a Countess 3 Automated Cell counter. We found that, for 10 pooled brains, this protocol produced 250,000-350,000 cells with a low (0.3-2%) cell death rate.

### Plating, media, and incubation

We coated surfaces for cell plating by incubating microwells in 30,000-70,000 Dalton Poly-L-lysine hydrobromide dissolved in 0.1 M Borate Buffer pH 8.4 (25 mM Borax, 0.1 M Boric Acid, 75 mM NaCl) at 28°C and 3% CO_2_ overnight (18-24 hours). The next day, coating solution was removed, surfaces were washed 5 times with copious sterile distilled water and left to dry for at least 1 hour before cells were plated. Cell suspensions were then pipetted onto the microwells’ surfaces to achieve a seeding density of 1000 cells per mm^2^, based on counted cell numbers, within 100 µl of seeding media which consisted of 10% Fetal Bovine Serum (FBS), 2% B-27 supplement, 1 mM GlutaMAX, 100 U Penicillin/Streptomycin in Neurobasal. Cells were allowed to attach for at least 1 hour before an additional 500 µl of seeding media was added.

For experiments with FBS, the media was not changed thereafter. To produce pure neuronal cultures, media changes were required to reduce the FBS concentration. As such, after 1 week of incubation in seeding media, 500 µl of culture media (2% ul B-27 supplement, 1 mM GlutaMAX, 100 U Penicillin/Streptomycin in Neurobasal) was added. For the rest of the culturing period, 50% of the media was changed on a weekly basis. Cells were incubated at 28°C and 3% CO_2_ until required for experimental purposes.

### Immunocytochemistry

We found that full aspiration of culture media resulted in the cells’ lifting off the microwell surface. To prevent this loss of cultured cells, all media changes, incubations, and washes were performed with a dead volume (100 µl) that was not removed. Cells were fixed using 4% PFA in 1 x Phosphate Buffered Saline (PBS) for 20 minutes at room temperature. Fixing solution was removed and the cells were washed 5 times using PBS. The cells were then incubated in blocking buffer (5% normal goat serum, 0.1% Triton X-100 in PBS) at room temperature for at least 40 minutes. Blocking buffer was removed and the cells were incubated in primary antibody solution (in blocking buffer) overnight at 4°C. The next day, cells were washed 5 times using PBS and then incubated in secondary antibody solution overnight at 4°C. Finally, cells were washed 5 times in PBS and stored in 70% glycerol. Antibodies used in this study include: rabbit anti-glutamate (1:500, Sigma-Aldrich, Cat #G6642), chicken anti-GABA (1:2000, Abcam, Cat #ab62669), mouse anti-SV2 (1:100, Developmental Studies Hybridoma Bank, Cat #SV2), rabbit anti-PSD-95 (1:2000, Abcam, Cat #ab18258), rat anti-Gephyrin (1:1000, Synaptic Systems, Cat #147208), Alex Fluor 635-conjugated goat anti-mouse IgG (1:2000, Invitrogen, Cat #A31574), Dylight 633-conjugated goat anti-rabbit IgG (1:2000, Invitrogen, Cat #35562), Alexa Fluor 568-conjugted goat anti-rabbit IgG (1:2000, Invitrogen, Cat #A11011), Dylight 488-conjugated goat anti-rat IgG (1:2000, Invitrogen, Cat #SA5-10018).

### Microscopy

For cellular analyses, cells were imaged on an Axio Observer Z1 inverted fluorescence microscope (Carl Zeiss) fitted with a full-enclosure incubator with CO_2_ and temperature control for live-cell imaging, an Axiocam 702 monochrome high-speed camera (Carl Zeiss), and a 20x/0.8 NA or 40x/0.6 NA Plan-Apochromat objective (Carl Zeiss). Image acquisition was performed using ZEN software.

For synaptic analyses, cells were imaged on a spinning-disk confocal system (Marianas; 3I, Inc.) consisting of an Axio Observer Z1 (Carl Zeiss) equipped with a CSU-W1 spinning-disk head (Yokogawa Corporation of America), ORCA-Flash4.0 v2 sCMOS camera (Hamamatsu Photonics), 63x/1.4 NA Plan-Apochromat objective. Image acquisition was performed using Slidebook 6.0.

### Image analysis

Analyses of cells based on phase contrast, DIC, or fluorescence were performed using ImageJ (Schneider et al., 2012). ROIs were generated by thresholding the Laplacian (FeatureJ plugin, smoothing scale of 4) of DAPI fluorescence images and applying ImageJ’s in-house particle detection analysis. The ROIs were used to measure antigen immunofluorescence intensity for individual cells, as well as the density of cells per culture and nuclear aspect ratio.

For the estimation of neuronal and non-neuronal cell number per culture, we used NeuN level and nuclear aspect ratio as distinguishing factors. Our rationales for choosing these factors were (1) NeuN is an known marker of neurons (Gusel’nikova and Korzhevskiy, 2015) and (2) we observed that many of the apparently non-neuronal cells *in vitro* appeared to either have elongated or irregularly-shaped nuclei (see Results). Classification of cells based on median anti-NeuN immunofluorescence intensity and nuclear aspect ratio were performed by applying Gaussian Mixed Model clustering in Python (Pedregosa, 2011). This approach produced three categories; (1) cells defined by high NeuN and low aspect ratio, (2) cells defined by low NeuN and high aspect ratio, and (3) cells that did not fit into the first two categories.

For the classification of neurons as GABAergic or glycinergic based on their immunofluorescence, we generated a threshold to identify outliers with high anti-GABA and anti-Glycine immunostaining intensity. We combined all measured modal staining intensities across all cultures (wild type) into a single population and extracted the first and third quartiles (Q1 and Q3), and the interquartile range. We defined an upper threshold based on three interquartile ranges above Q3 and applied the threshold to each culture (across all genotypes) separately such that any neuron with values above this would be considered GABAergic or glycinergic. We expressed the percentage of GABAergic or glycinergic neurons on a per-culture basis and used these values for further analysis. To find the percentage of *gad1b*-positive neurons we manually counted the percentage of fluorescent neurons across 3 separate cultures collected from embryos harboring *gad1b:GAL4* (Förster et al., 2017) and UAS:GCaMP6f.

Analysis of synapses based on confocal stacks was performed using Imaris 9.6 (Bitplane). Immunoreactivity against SV2 was used to label synaptic vesicles, as described previously (Du et al., 2018). We decided that rendering a surface (rather than spheres) based on SV2 immunostaining intensity was the most appropriate approach for representing presynaptic regions in our cultures because of how SV2 is distributed throughout boutons. We chose an intensity threshold to render the surfaces by closely matching the underlying immunostaining of three separate pilot confocal stacks and averaging the threshold across all three. This average was ultimately applied to all stacks for formal analysis. We ensured that the threshold for all three pilot stacks did not produce extraneous surfaces, which can occur in high fluorescence intensity regions if the threshold is too low. The consequence of choosing a threshold based on this approach is that less intense SV2 regions do not get rendered as a surface. We operated under the assumption that much of the low intensity staining against SV2 represented synaptic vesicles not specific to active synaptic regions and so should not be rendered into the final reconstruction. It is possible that some synapses, especially small ones, may not be accounted for when taking this approach. We took a similar approach to rendering postsynaptic sites based on immunoreactivity against PSD-95 and Gephyrin (Gphn). Given the specificity of the labelling for PSD-95 and Gphn, we could observe clear puncta, and thus chose to render them as spheres rather than surfaces. As with the SV2 approach, we selected intensity thresholds across three pilot stacks based on the criteria that they did not produce extraneously rendered spheres (based on background) and averaged the threshold values for application to formal analysis. Once all three channels had been rendered, we applied an exclusion criterion to PSD-95- and Gphn-based puncta based on their distance from SV2 surfaces (300 nm). Once all but the closest puncta were excluded, we counted those remaining and normalized the value to the total SV2-based volume to account for inter-culture and inter-stack variability in synaptic density.

### Neuronal activity analysis

HuC:GCaMP6s fluorescence activity was recorded as CZI files before being converted to TIF files using ImageJ (Schneider et al., 2012). We then used the calcium imaging package, Suite2p (Version 0.10.3), to identify regions of interest (ROIs) and extract fluorescence signals from the ROIs. The default Suite2p cell classifier was customized to our cell culture images using manually labelled ROI data. All ROIs were then classified as cells or not cells using this updated classifier (Fig. S1A). The same parameters for Suite2p were used across recordings from cultures at days *in vitro* (DIV)1, 7, 14 and 21 (n=1). Across all recordings, 589 ROIs (DIV 1=115, DIV 7=154, DIV 17=133, DIV 21=187) were detected and extracted.

For the analysis of ROI activity traces, we used MATLAB (R2021a) and computed ΔF/F as has been previously described (Vanwalleghem et al., 2021, Akerboom et al., 2012). ROIs with ΔF/F activity >0.3 were selected and visualized for each recording (Fig. S1B-D).

### Analysis of Bulk RNA-seq Datasets

Differentially expressed genes from a bulk RNA-sequencing dataset of *scn1lab^Δ44^* mutants from Weinschutz Mendes et al. (2023) were analyzed. RNA extraction and sequencing methods are described in Weinschutz Mendes et al. (2023). Briefly, total RNA was extracted from isolated larval heads with the eyes removed (n=3-4 heads per genotype, 1 head per sample) from sibling or cousin-matched *scn1lab^+/+^*, *scn1lab^Δ44/+^,* and *scn1lab^Δ44/Δ44^* larvae at 6 dpf. The RNA library was sequenced at 25 million reads/sample using NovaSeq 6000 (Illumina) (Yale Center for Genome Analysis). Raw reads are accessible at Gene Expression Omnibus (Accession # GEO: GSE205578). Low quality reads were trimmed and adaptor contamination removed using Trim Galore (v0.5.0) and trimmed reads were mapped to GRCz11 *Danio rerio* reference genome using HISAT2 (Kim et al., 2019). Gene expression levels were quantified using StringTie (v1.3.3b) (Pertea et al., 2015) with gene models (release 103) from Ensembl. Differential expression analysis was conducted using DESeq2 (v1.22.1) (Love et al., 2014) and results are accessible at Gene Expression Omnibus (Accession # GEO: GSE205578). Significantly differentially expressed genes were determined by p-adjusted value of <0.05 based on DESeq2 analysis. Upregulated or downregulated gene expression was determined by the sign of the log2 Fold change, where positive sign is upregulated and negative sign is downregulated. Normalized counts were ascertained by DESeq2 analysis and significant differences in counts between *scn1lab^Δ44/Δ44^* and *scn1lab^+/+^* were determined by p-adjusted value of <0.05.

### Statistics

A combination of MATLAB, Python 3.7, and Graphpad Prism version 9 for Windows (GraphPad Software) were used to test for statistically significant differences between genotypes and generate graphs. For all data, we used a Shapiro-Wilk test to check data for normality. Where data followed a normal distribution, we used a one-way analysis of variance (ANOVA) with Holm-Šídák’s post hoc test for multiple comparisons, otherwise we used a Kruskal-Wallis test with Dunn’s post hoc test for multiple comparisons. Significance was determined at *p* < 0.05.

## Results

### Long term primary culture of embryonic zebrafish neurons

With the aim of optimizing primary neuronal cultures from zebrafish brain tissue, we tested the effects of a range of variables on *in vitro* neuronal population quality and longevity. We found that zebrafish brain cell cultures were particularly sensitive to the age of the animal from which cells were harvested, the method of polymer coating, and the composition and volume of the culture media used. We attempted to culture brain cells from animals at different ages from 1-6 dpf and found a developmental window from which cell survival was maximized. Specifically, animals between 30 and 36 hpf produced robust cultures with the lowest rates of cell death.

Cells were highly resilient to mechanical trituration following enzymatic dissociation, and as such, we were able to generate homogenous single cell suspensions by triturating rapidly and extensively. Polymer coating worked best when performed in high pH Borate Buffer, with a small but noticeable improvement when using Poly-L-lysine (30,000-70,000 Da) instead of Poly-DL-ornithine (3,000-15,000 Da) or Poly-D-lysine (70,000-150,000 Da). We found that cell attachment and survival during the first week of the culture was sensitive to the presence of FBS and the volume of media in which the cells were grown. Maintaining a relatively low volume of media with 10% FBS produced optimal survival, so we only added FBS-free culture media after day *in vitro* (DIV) 7, followed by weekly half media changes. Using this protocol, we consistently produced high quality neuronal cultures that visibly formed networks comparable to those observed when using well established cell lines or mammalian embryonic brain tissue (Fig. 1A-D). We were able to maintain cultures for over 100 days.

**Figure 1.**
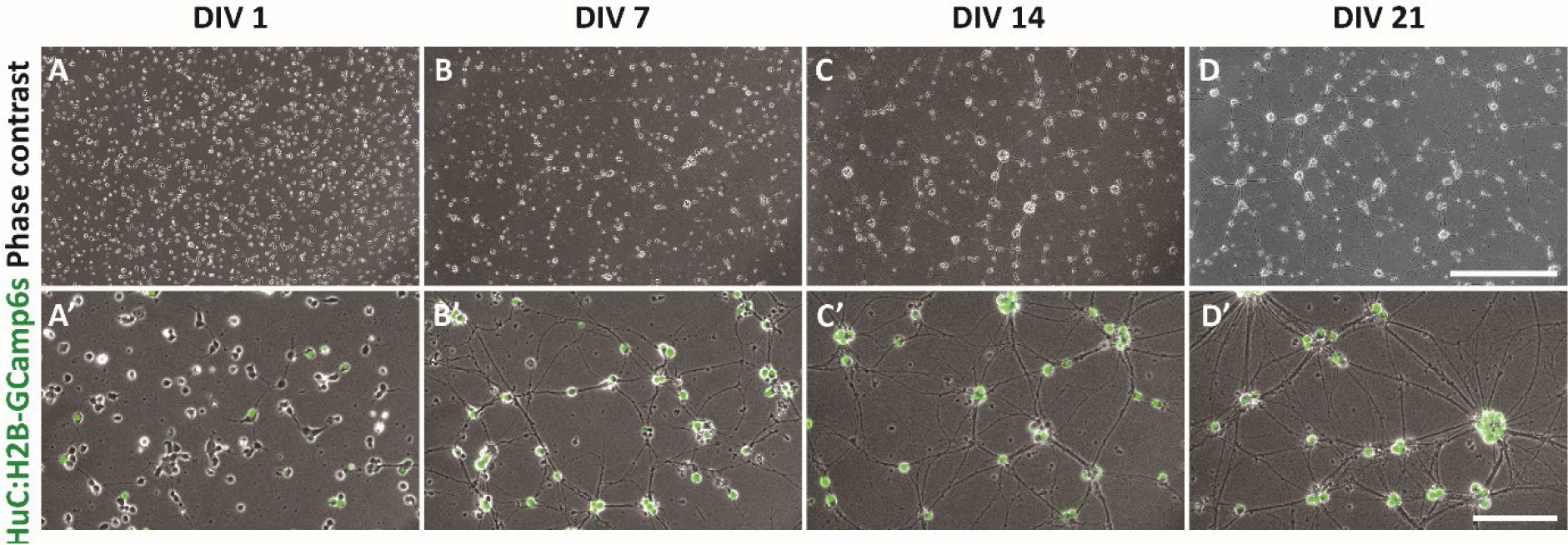
Overview of pure primary zebrafish neuronal culture. (A-D’) Representative images of pure neuronal cultures derived from 30-36 hpf zebrafish embryos harboring *HuC:GCaMP6s*. Phase contrast alone (top) and merged with *HuC:GCaMP6s* fluorescence (bottom) at 1 (A, A’), 7 (B, B’), 14 (C, C’) or 21 (D, D’) days *in vitro* (DIV). Scale bars, 500 μm (top), 100 μm (bottom).

By virtue of the brain tissue purity and sparsity of cells overall, we could easily document major developmental milestones of the neuronal networks as they matured. Doing so from animals that expressed a nuclear GCaMP6s reporter under control of HuC (*elavl3*) allowed us to identify neurons (Kim et al., 1996) and observe their activity. Within the first week, neurons predominantly grew projections and established initial contacts through thin neurites (Fig. 1A’). Although abundant initially, HuC-negative non-neuronal cells rapidly died leaving mostly neurons by DIV 7 (Fig. 1B’). In the second week, presumptive synaptic connections began to appear and interconnecting cellular processes thickened, likely due to projection and fasciculation of additional neurites along major tracks (Fig. 1C’). Furthermore, thin neurites projected into unoccupied spaces and swellings (reminiscent of dendritic spines or presynaptic boutons) started to appear along neurite tracks (Fig. 1C’). In the third week, the thickening of fascicles and emergence of swellings continued until these processes reached a plateau, after which the cultures’ morphology stabilized (Fig. 1D’). Fluorescence imaging of GCaMP6s revealed that neurons displayed activity from DIV 1 through at least DIV 21 (Fig. S1B), which was the last time point at which we made fluorescence observations.

**Figure S1.**
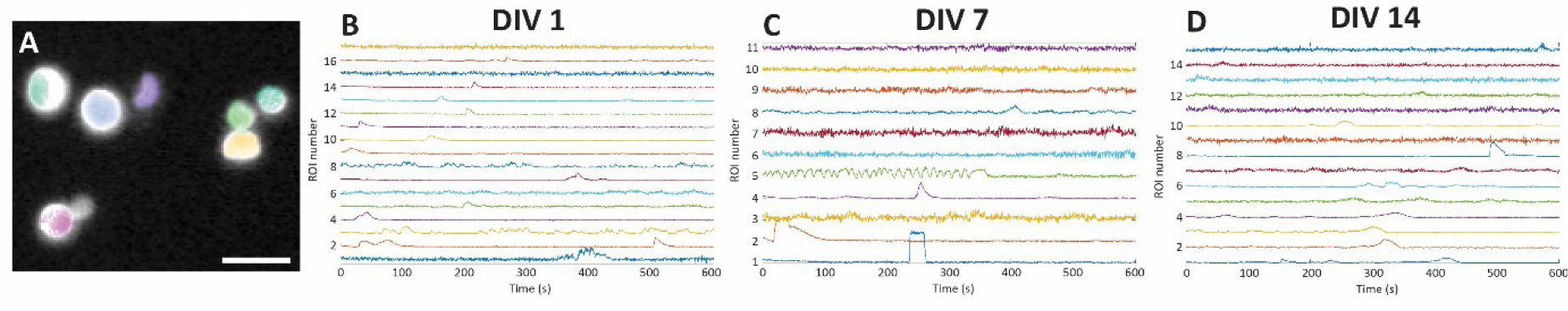
Calcium imaging of neuronal activity *in vitro*. (A) Representative images of detected ROIs overlayed on GCaMP6s fluorescence of neuronal nuclei. Cultures from fish harboring *HuC:H2B-GCaMP6s* were imaged to extract information about neuronal activity. Suite2p was used to detect ROIs associated with individual neuronal nuclei and extract calcium traces. Scale bar 20 μm. (B-D) Nuclear calcium traces (ΔF/F) extracted from cultures demonstrate that neurons are active *in vitro*. Heterogenous levels of activity were observed at DIV 1 (B), DIV 7 (C) and DIV 14 (D).

### Assessment of *in vitro* cell type composition

To begin characterizing the cultures produced by our protocol, we sought to identify the major cell types populating them. We expected high neuronal purity given our specific dissection of brain tissue but were also interested in the presence of surviving glia. The 30-36 hpf window is a period of intense neurogenesis (Wullimann and Knipp, 2000), and therefore many multipotent cells were likely to be present in our whole-brain dissociates. Our choice of growing cultures in FBS for at least the first week of culture is relevant to the eventual cellular composition of the cultures, since the factors present in FBS typically promote survival and differentiation of non-neuronal cells (Hashimoto et al., 2000, Kinikoglu et al., 2014). To explore the effect of FBS, we cultured cells under two separate conditions: one where we reduced the FBS concentration via weekly half FBS-free media changes and one with no media changes at all (10% FBS present from beginning to end). To validate the broader utility of our protocol we also investigated the effects of these different culturing conditions on cells from *scn1lab* mutant embryos and their wild type siblings. Mutations in *scn1lab* produce well-documented phenotypes in zebrafish (Banerji et al., 2021, Baraban et al., 2013, Brenet et al., 2019, Schoonheim et al., 2010, Tiraboschi et al., 2020, Weinschutz Mendes et al., 2023) and as such, it is a good candidate for testing the sensitivity of our methodology for detecting neuropathological phenotypes *in vitro*.

Under conditions without FBS, we found only presumptive neurons, on the basis that all cells had small round somata, contained acetylated tubulin, and stained positive for NeuN, which is a well-established marker of mature neurons (Gusel’nikova and Korzhevskiy, 2015). This interpretation was also supported by cultures from *Huc:H2B-GCaMP6s* transgenic animals, with all observed cells expressing GCaMP6s (Figs 1D’) off the neuronal HuC (*elavl3*) promoter (Kim et al., 1996). With FBS, however, we found several other cell types surviving alongside neurons. These cells were GFAP-negative, negative for acetylated tubulin, and stained poorly for NeuN, implying they were not neurons or astroglia. They exhibited a range of morphologies including flat, oblong, spindly, or stellate, indicating a diversity of cell types (Fig. S2A-D). Furthermore, some formed clear associations with neurons while others appeared not to interact with other cells (Fig.S2A-D).

Gaussian Mixed Model clustering using NeuN staining intensity and nuclear aspect ratio allowed us to categorize cells as neuronal (defined by high NeuN and low aspect ratio) or non-neuronal (defined by low NeuN and high aspect ratio). Not all cells fell into these two categories, however, resulting in a third non-specific category of unclassified cells. Quantification of the categories in our cultures with FBS revealed that on average, 78.4% of cells could be classified as neurons, 4.4% as non-neuronal and 17.2% could not be categorized (n=3 independent cultures, DIV 27-31). Cultures produced from homozygous *scn1lab* mutant animals looked strikingly different compared to those from their wild type siblings (Fig. 2A, B). These differences appeared to be due to an increased number of cells that stained weakly for NeuN and acetylated tubulin (Fig. 2C-C’’, D-D’’). Application of Gaussian Mixed Model clustering revealed clear differences in the proportions of cell types populating *scn1lab* mutant cultures compared to wild type (Fig. 2E-G). Quantification of cell density revealed that *scn1lab* heterozygous and homozygous mutant cultures had 64% and 88% more cells than wild type cultures, respectively (Fig. 2H). The increase in total cell density could not be attributed to a change in the number of neurons across the three genotypes (Fig. 2I). Instead, the average density of non-neuronal cells in *scn1lab*^−/−^ cultures was approximately 15 times higher than wild type, representing a statistically significant difference (Fig. 2J). As a result of the specific increase in non-neuronal cells, the proportion of these cells was significantly higher in *scn1lab*^−/−^ cultures, 34% on average (up from 4.4% in wild type, Fig. 2K). Cultures generated from heterozygous *scn1lab*^−/+^ brains showed an intermediate phenotype, but the density of non-neuronal cells was not significantly different from the those in *scn1lab*^+/+^ or *scn1lab*^−/−^ cultures. Because there were no clear or significant differences in the density of neuronal cells across genotypes, these data demonstrate that the phenotype is most likely attributable to altered proliferation or differentiation of non-neuronal cells after plating. This is because the seeding density was kept constant across all three genotypes. If the increase in non-neuronal cells were due to a higher proportion of these being present at the time of plating, the number of neurons would be expected to be lower. Our data thus suggest that *scn1lab* suppresses cell proliferation of at least one multipotent population of cells present in 30-36 hpf embryonic brains that differentiates predominantly into non-neuronal cells.

**Figure 2.**
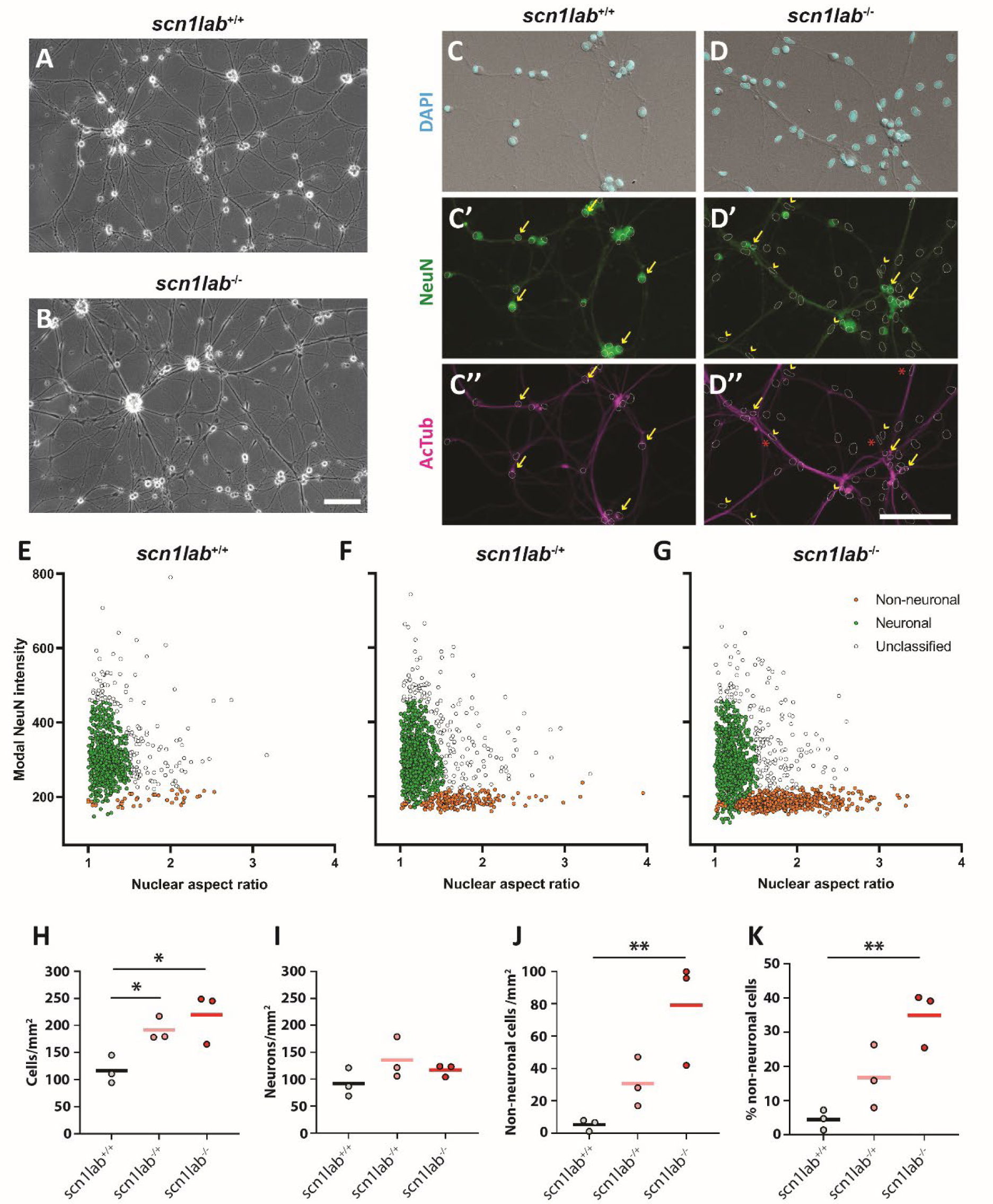
Prevalence of non-neuronal cells in mixed cultures. (A-B) Representative phase contrast images of cultures derived from (A) DIV 27 wild type (*scn1lab*^+/+^) and (B) DIV 29 *scn1lab* mutant homozygote (*scn1lab^−/−^*) animals. Scale bar, 100 µm. (C-D’’) Fluorescence imaging of wild type and *scn1lab* mutant cultures for classification of major cell types. Nuclei are revealed by labelling of DAPI (blue, C-D) and are used to generate ROIs for further analysis (dotted lines). Cells with high immunoreactivity against NeuN (yellow arrows, C’-D’) often appear to have round nuclei and are likely neurons while cells with low immunoreactivity against NeuN (yellow arrowheads, D’) often have irregular nuclei and are likely non-neuronal. NeuN-positive cells often appear to contain acetylated tubulin (yellow arrows, C’’-D’’) whereas NeuN-negative cells do not (yellow arrowheads, D’’). Further, some NeuN-negative cells appear closely associated with tubulin-rich axons and are often found nestled within axon bundles (red asterisks, D’’). Scale bar, 100 µm. (E-G) Gaussian Mixed Model clustering based on NeuN modal intensity and nuclear aspect ratio produces three categories of cells: non-neuronal (orange dots), neuronal (green dots) and non-specific/unclassified (white dots). (H-K) Quantification of cell numbers based on clustering results. Wild type are shown in grey, *scn1lab* mutant heterozygotes are shown in pink and *scn1lab* mutant homozygotes are shown in red. Each data point represents the average per culture, where a culture is defined by at least 10 pooled embryonic brains in an independent dissociate. Horizontal bars show average per genotype. Shapiro-Wilk test revealed all groups were normally distributed. Groups tested for statistically significant differences using Ordinary one-way ANOVA with Holm-Šídák’s multiple comparisons, * indicates *p*<0.05, ** indicates *p*<0.01.

**Figure S2.**
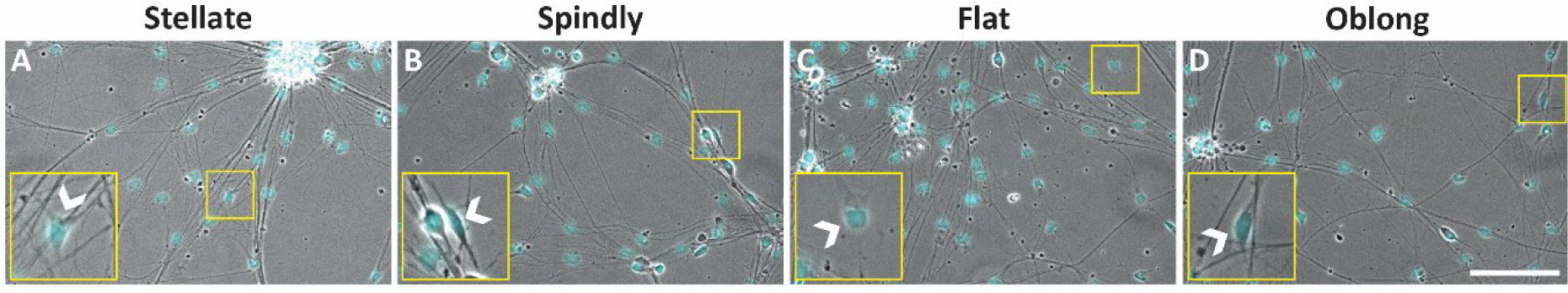
Morphology and molecular markers of non-neuronal cells in mixed cultures. (A-D) Representative DAPI-stained phase contrast images of DIV 100 cultures derived from wild type animals. Arrowheads in insets indicate different types of non-neuronal cells with diverse morphologies including (A) stellate, (B) spindly, (C) flat, and (D) oblong. Scale bar, 100 μm.

### Neuronal neurotransmitter subtypes in primary culture

The neurotransmitter subtype composition of zebrafish primary neuron cultures has yet to be explored in depth and has thus far been explored in a targeted way for neuromodulators (Patel et al., 2019; Tapanes-Castillo et al., 2014). One study reported over 90% of cultured zebrafish neurons expressed tryptophan hydroxylase 2 (Tapanes-Castillo et al., 2014), while another found that 85% of cultured neurons were positive for serotonin while all were negative for tyrosine hydroxylase and choline acetyltransferase (Patel et al., 2019). We set out to measure the main neurotransmitters utilized by excitatory and inhibitory neurons in our pure neuronal cultures with the goal of ensuring that our protocol produces a heterogenous population of cells roughly matching those in the brain.

To assess neurotransmitter type, we used antibodies raised against glutamate, GABA, and glycine, which have previously been used to identify neuronal subtypes in zebrafish (Higashijima et al., 2004, Martin et al., 1998, Pedroni and Ampatzis, 2019). We found that anti-glutamate immunoreactivity produced a poor readout for identifying neuronal subtypes, with widespread and homogeneous staining intensity across all cell bodies (Fig. 3B). This was despite clear indications of the antibodies’ functionally labelling glutamate, given we often observed bright puncta along axon fascicles consistent with glutamate-rich excitatory synaptic terminals (Fig. 3B, arrowheads). Antibodies against GABA and glycine, on the other hand, produced discriminable and quantifiable differences in staining intensity across different neuronal somata, indicating that they were appropriate labels for neurotransmitter subtypes (Fig. 3C, D). Some neurons that stained strongly for GABA had weak immunoreactivity against glycine and vice-versa (Fig. 3C, D, asterisks), consistent with *in vivo* observations by 4-5 dpf (Higashijima et al., 2004). In order to estimate the percentage of GABAergic and glycinergic neurons, we searched for cells with a high-intensity GABA and glycine immunoreactivity, respectively (Fig. 3F, G). By setting a strict outlier-oriented threshold, our analyses determined that roughly 0.4% of neurons were GABAergic, and 1.4% were glycinergic (n=7 cultures, DIV 32-35), implying that the majority were glutamatergic. We also grew cultures from double transgenic animals carrying *gad1b:Gal4* (Förster et al., 2017) and *UAS:GCaMP6f* as an independent approach for estimating the prevalence of GABAergic neurons (Fig. 3E). In these cultures, we found just over 0.4% of neurons expressing GCaMP6f (n=3 cultures, DIV 20), closely matching the value that we found using anti-GABA antibodies. Thus, our method produces a heterogenous population of neuronal subtypes with a strong bias towards excitatory neurons. When we quantified the number of inhibitory neurons in cultures grown from *scn1lab*^−/+^ and *scn1lab*^−/−^ embryos versus wild type siblings, we found that they had on average, roughly an order of magnitude fewer neurons classified as GABAergic (Fig. 3F). This difference was not statistically significant despite the trend, likely due to the low abundance of strongly GABA-immunopositive neurons overall (n=7 cultures, DIV 32-35). Importantly, this observation is consistent with known effects of *scn1lab* KO, which results in fewer GABAergic neurons *in vivo* (Tiraboschi et al., 2020). There was no such trend for neurons classified as glycinergic (Fig. 3G), suggesting specificity towards GABAergic cells.

**Figure 3.**
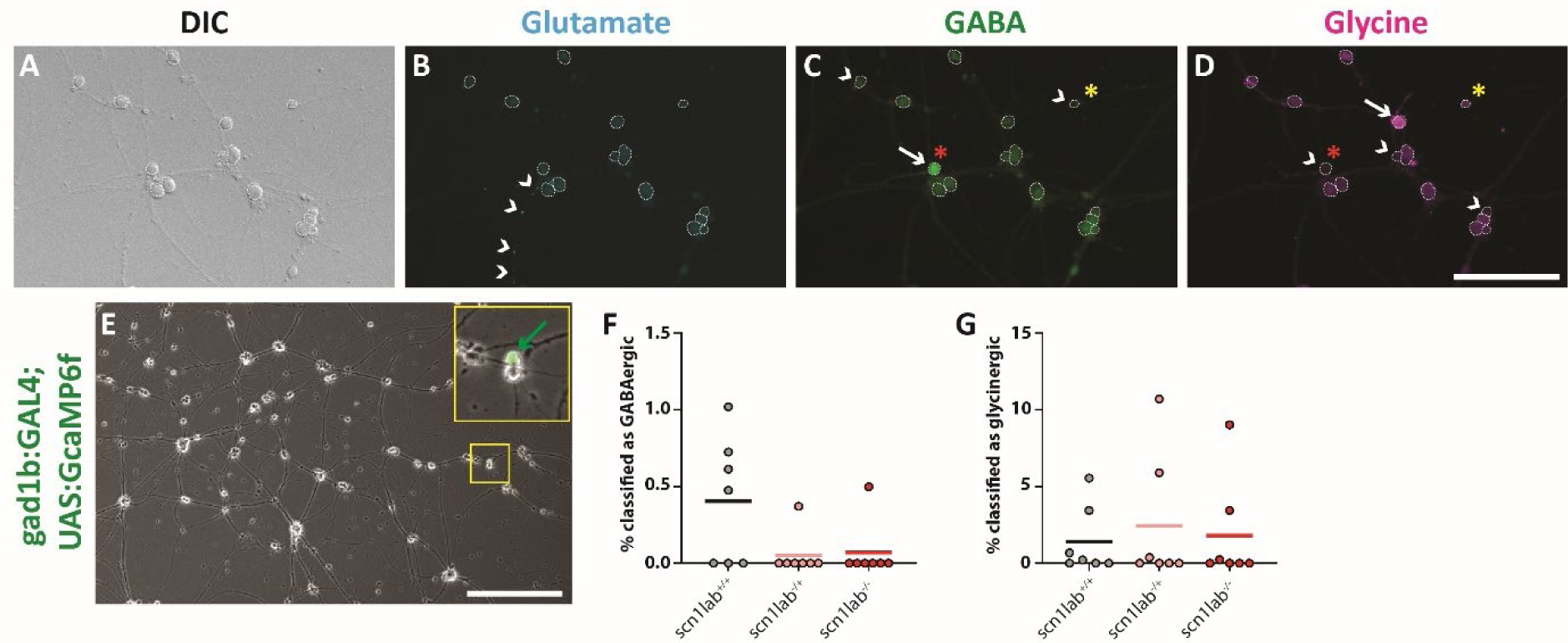
Neurotransmitter subtypes in primary cultures. (A-D) DIC image (A) and immunocytochemical labelling of glutamate (B), GABA (C) and glycine (D) from a representative 29 DIV pure neuronal culture. White dotted circles outline cell bodies from DIC image (A). White arrowheads in (B) show punctate glutamate immunoreactivity, indicating presumptive glutamate-rich excitatory synaptic puncta. White arrows in (C) and (D) show cells with strong staining against GABA and glycine respectively, whereas white arrowheads show cells with weak staining. Red asterisk denotes a cell with high GABA and low glycine. Yellow asterisk denotes a cell with relatively high glycine and low GABA. Scale bar, 100 µm. (E) Representative phase contrast image of DIV 20 pure neuronal culture derived from animals harboring *gad1b:GAL4* and *UAS:GCaMP6s*. GCaMP6s-expressing neuron (*gad1b*-positive) indicated with green arrow (yellow inset) Scale bar, 250 µm. (F, G) Quantification of GABAergic (F) and glycinergic (G) neuron percentages in pure neuronal cultures acquired by applying an outlier threshold on fluorescence intensity of anti-GABA and anti-glycine staining, respectively. Cultures were grown from wild type (black), *scn1lab* heterozygous mutants (pink) and *scn1lab* homozygous mutant (red) siblings. Each data point represents the average per culture, where a culture is defined by at least 10 pooled brains in an independent dissociate. Horizontal bars show average per genotype. Shapiro-Wilk test revealed all groups were not distributed normally. Groups tested for statistically significant differences using Kruskal-Wallis test with Dunn’s multiple comparisons.

### Analysis of synaptic connectivity

One key feature of functional neuronal networks is the establishment and maintenance of mature synaptic connections. Synaptic markers have not been applied to zebrafish primary brain cell cultures before, and as such, it is difficult to ascertain whether these cultures form functional networks that could model brain activity. With this in mind, we set out to measure whether our cultured neurons bore the structural hallmarks of mature synapses. We designed an immunostaining strategy for labelling mature synapses by defining maturity as obligate opposition of presynaptic (SV2, Fig. 4A-A’’) and postsynaptic (Gephyrin, Fig. 4B-B’’ and PSD-95, Fig. 4C-C’’) markers, whereby the postsynaptic markers had accumulated into quantifiable puncta (see Methods). Defining synapses this way, we found evidence for abundant synaptic connectivity in our pure neuronal cultures (Fig. 4D-F, n=7 cultures, DIV 32-35). We found that there was a consistently higher number of excitatory connections than inhibitory ones, with the PSD-95:Gphn ratio ranging from 1.5:1 to 3:1 in different cultures (Fig. 4G). This predominance of excitatory synaptic transmission is consistent with our previous findings that a minority of neurons in these cultures are inhibitory. Cultures sourced from wild type, *scn1lab*^−/+^ and *scn1lab*^−/−^ animals showed similar numbers of PSD-95 puncta and Gphn puncta, and similar values for the PSD-95:Gphn ratio (Fig. 4E, F, G, n=7, DIV 32-35).

**Figure 4.**
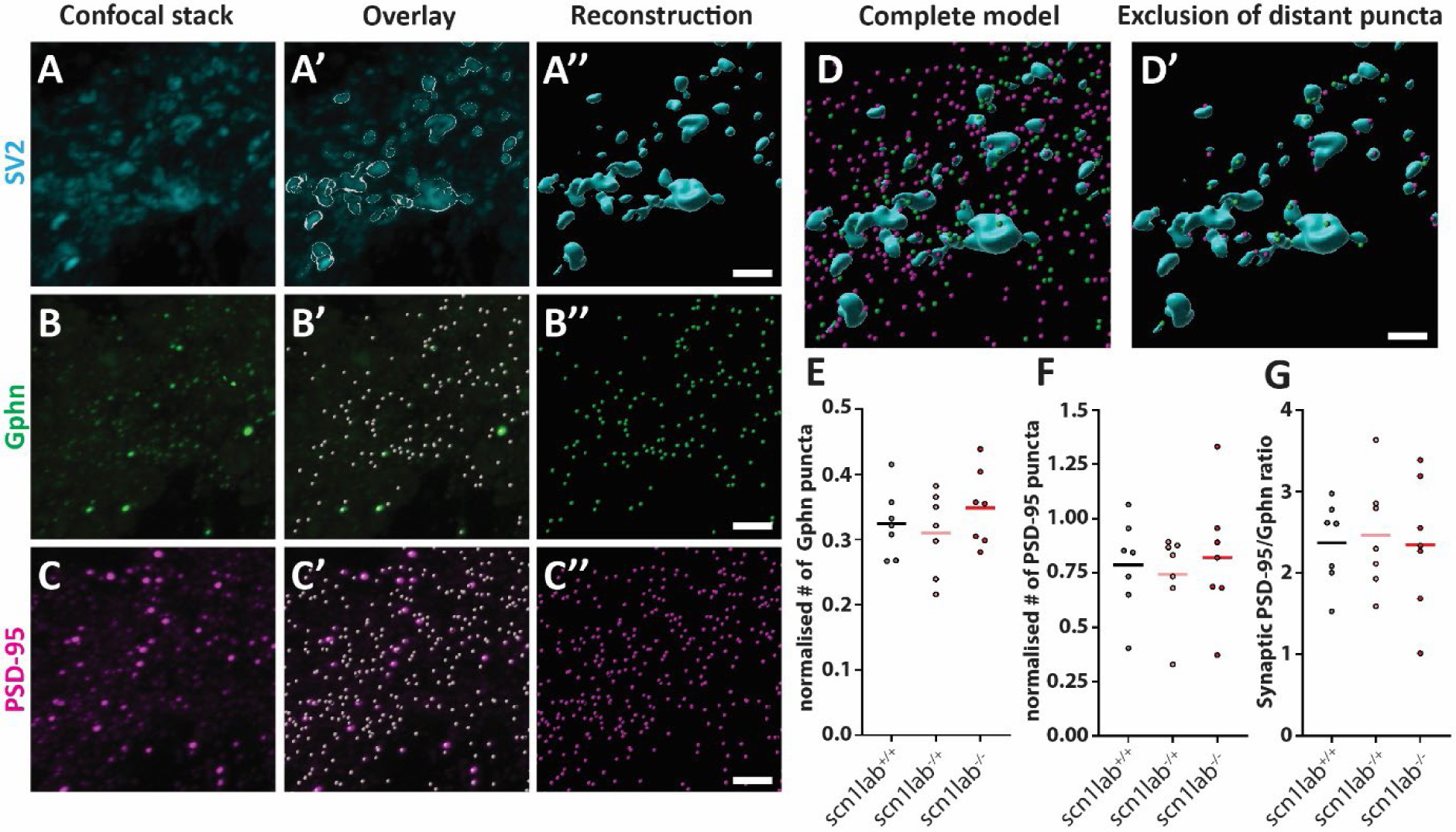
Assessment of excitatory and inhibitory synapses in pure neuronal cultures. (A-C’’) Representative images from Imaris 3D projected confocal stack of synapses in pure neuronal cultures. Immunoreactivity against synaptic markers was used to generate 3D reconstructions, upon which further analyses were based. (A-C’’) Progression of rendering shown from initial immunofluorescence, to overlay with 3D render, and lastly, isolated reconstruction for SV2 surfaces (A-A’’), Gephyrin spheres (B-B’’) and PSD-95 spheres (C-C’’). (D-D’) Process of sphere exclusion based on proximity to SV2 surfaces (300 nm) transforms full reconstruction (D) to synaptic-only reconstruction (D’), where SV2 surfaces are blue, Gphn spheres are green and PSD-95 spheres are magenta. Scale bars, 3 μm. (E-G) Quantification of the number of Gphn puncta normalized to SV2 volume (E), the number of PSD-95 puncta normalized to SV2 volume (F), and the ratio of PSD-95:Gphn puncta per culture (G). Cultures were grown from wild type (black), *scn1lab* heterozygous animals (pink) and *scn1lab* homozygous mutant (red) siblings. Each data point represents the average per culture, where a culture is defined by at least 10 pooled brains in an independent dissociate. Horizontal bars show average per genotype. Shapiro-Wilk test revealed all groups were normally distributed. Groups tested for statistically significant differences using Ordinary one-way ANOVA with Holm-Šídák’s multiple comparisons.

### Transcriptomic validation of predicted *in vitro scn1lab* phenotypes

Results from our primary neuron cultures pointed toward particular developmental phenotypes. Specifically, our cultures indicated increased proliferation in cultures derived from *scn1lab* mutants (Fig. 2), a higher proportion of non-neuronal cells in mutant-derived cultures (Fig. 2), and a trend toward fewer GABAergic neurons in cultures derived from mutants (Fig. 3F). One of the promising uses for primary neuronal culture is to identify molecular or physiological phenotypes in a tractable culture, and in so doing, provide leads for targeted *in vivo* experiments for the same effects. With leads from our culture experiments, we looked for corresponding phenotypes in *scn1lab* mutant animals.

Using a previously published dataset of *scn1lab* bulk transcriptomic data from 6 dpf animals (Weinschutz Mendes et al., 2023), we looked for transcriptional profiles that could provide support for our observations of altered cell-type abundance. We found that the excitatory and inhibitory neuronal transcripts, *slc17a6b* and *gad1b*, respectively, were both downregulated in homozygous *scn1lab* mutants relative to wild type, which suggests decreased relative abundance of these cell types (Figure 5A). This mirrored our finding of altered cell type composition *in vitro* due to enhanced proliferation of non-neuronal cells (Figure 2K). In agreement with this, we found a significant upregulation of *tyrp1a* and *tyrp1b* transcripts in *scn1lab* mutants (Figure 5A), indicative of enhanced abundance of non-neuronal melanin-expressing cells. There was no difference in the number of glial transcripts *mbpa*, *mbpb*, *mag*, *S100b* and *gfap* (Figure 5C), suggesting that the increase in non-neuronal cells is specific to a lineage that can differentiate into melanocytes (Dawes and Kelsh, 2021). It should be noted that the tissue used for transcriptomic analysis included brain and dermal tissue from *scn1lab* larvae, which are notably hyperpigmented. It is therefore possible that the signals indicating increased prevalence of melanin-expressing cells have an origin in dermal tissue and not the brain. Importantly, this result is consistent with our *in vitro* observations because melanocytes are derived from multipotent neural crest cells. Moreover, our analyses here are consistent with the enrichment of neuronal, GABAergic and glutamatergic markers among downregulated genes in *scn1lab* mutants at 6 dpf and an increase in the number of mitotic, PH3-positive, cells at 3 dpf (Weinschutz Mendes et al., 2023). Combined, these results show that our *in vitro* cultures effectively recapitulate changes that are detectable in intact organisms and provide confidence that they can be used to meaningfully explore neurobiological gene function.

**Figure 5.**
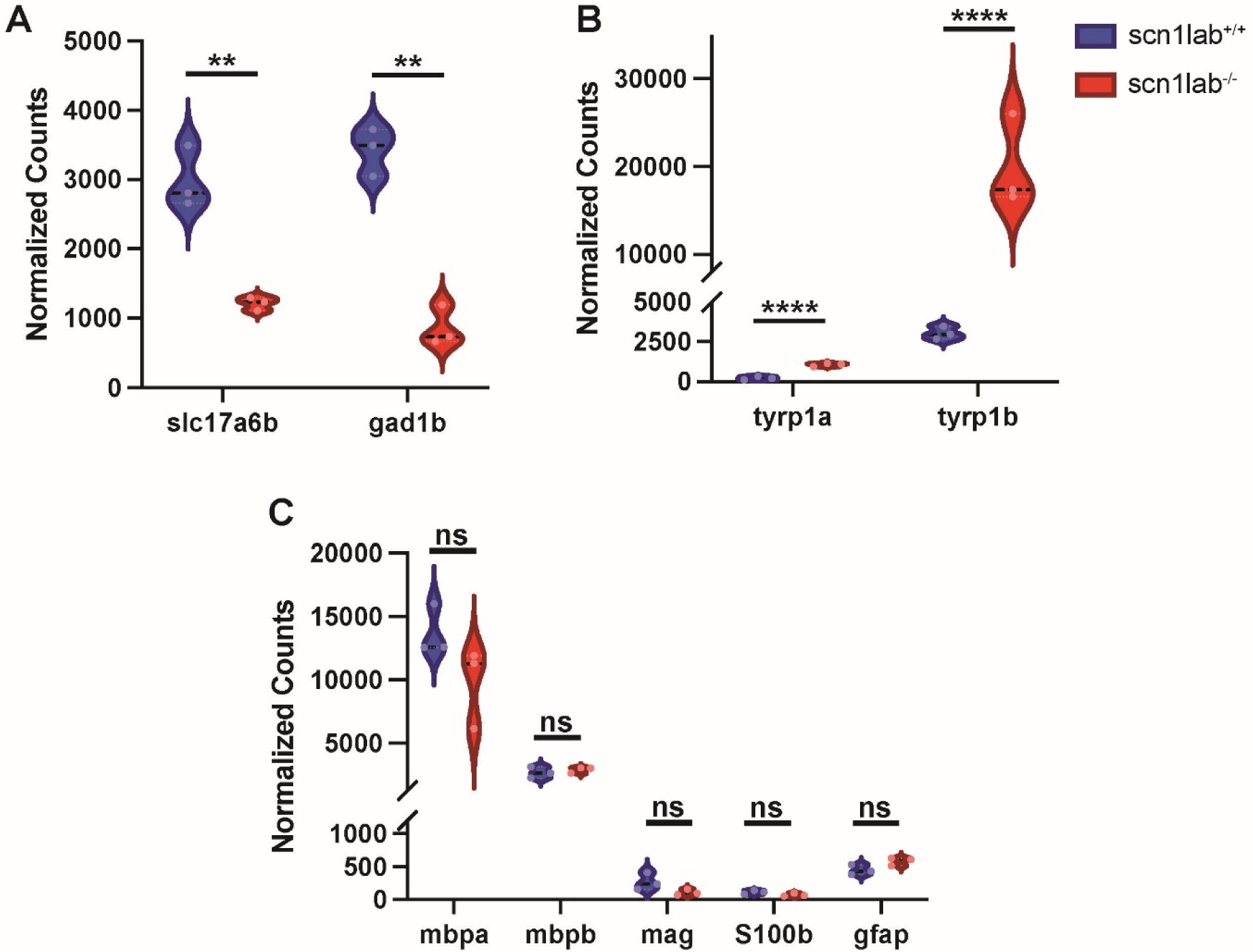
RNA-seq identifies dysregulation of neuronal and non-neuronal transcripts in *scn1lab* mutants. Violin plots showing normalized counts for select neuronal and non-neuronal genes in *scn1lab*Δ*44/*Δ*44* and *scn1lab+/+* fish at 6 dpf. **(A)** Downregulation of glutamatergic and GABAergic markers; **(B)** upregulation of melanocyte markers; and **(C)** no significant difference in glial cell markers in *scn1lab*Δ*44/*Δ*44* compared to *scn1lab+/+* fish. ****p-adjusted<3×10-5; **p-adjusted<0.005.

## Discussion

Current protocols for generating primary zebrafish neuron cultures have yet to reach the high purity and longevity typically associated with well-established cell lines and mammalian primary neuron cultures (Ray et al., 1993, Vicario-Abejón, 2004, Wiatrak et al., 2020). Here, we report a protocol that produces pure primary zebrafish neuron cultures that can be maintained for long periods of time. We were able to culture primary neurons for over 100 days, indicating comparable longevity to primary rodent hippocampal cultures that can grow for weeks to months (Ray et al., 1993; Vicario-Abejón, 2004). The neurons in our cultures formed visible structural networks and displayed activity measurable with GCaMP6s (Fig. S1). Thus, we believe the protocol we present here represents the current best practice for generating and maintaining primary neuronal cultures from zebrafish.

We found several key factors influencing the quality and longevity of our cultures. Likely the most important of these was the age of the embryo from which we sourced dissected brains. Neurons cultured from 30-36 hpf embryos had the highest level of long-term survival, indicating a uniqueness to this developmental period. It is not surprising that there should exist an optimal developmental window from which to harvest brain cells for culture. Protocols for culturing hippocampal neurons in rodents describe a developmental window around embryonic day 16-18 during which the health of subsequently cultured neurons is maximized (Ray et al., 1993; Vicario-Abejón, 2004). Interestingly, 30-36 hpf in zebrafish is a window that lies between two main waves of developmental neurogenesis: primary neurogenesis, which occurs before 24 hpf and secondary neurogenesis, which occurs after 48 hpf (Kimmel, 1993, Mueller and Wullimann, 2003). As such 30-36 hpf may represent a window during which most cells have established a neuronal identity but have not yet formed such extensive interconnections that enzymatic and mechanical dissociation produces fatal injury.

We characterized several aspects of our cultures with the goal of demonstrating that they formed synaptic networks comprising heterotypic neurons, akin to those observed in the brain. We found that having 10% FBS throughout the entire duration of the cultures’ growth resulted in the survival of cells that stained poorly for NeuN and acetylated tubulin (Fig. 2). We estimated that by DIV 27-31 these cells represented approximately 4.4% of the population and displayed a range of morphologies (Fig. S2), indicating a diversity of cell types. Removing FBS via half-weekly media changes largely abolished these cells and resulted in pure neuronal cultures (Fig. 1).

Although we have yet to find a marker that clearly identifies the cells present in the FBS-containing cultures, some had obvious morphological traits that suggested their identity. One type, for example, exclusively formed tight interactions with axon bundles and had elongated nuclei (Fig. 2D-D’’). These were highly reminiscent of Schwann cells *in vivo*, which similarly possess elongated nuclei and are located within axon bundles (Xiao et al., 2015). Others had characteristics of neural crest cells (NCCs) in that they were flat and stellate (Fig. S2), appearing similar to those previously purified and cultured from zebrafish embryos (Kinikoglu et al., 2014). Survival of cultured zebrafish NCCs was found to be highly sensitive to the presence of FBS (Kinikoglu et al., 2014), so their survival in our FBS-containing cultures is consistent with their known properties. The survival of NCCs provides an explanation for the range of diverse cells we observed when culturing with FBS, given their multipotency. For example, the entire Schwann cell lineage is descended from NCCs (Jessen and Mirsky, 2019).

We used an antibody labelling strategy to show the percentage of GABAergic and glycinergic cells in our pure neuronal cultures between DIV 32-35. We found that we could identify GABAergic and glycinergic cells by selecting strongly immunopositive cells (using antibodies against GABA and glycine) via a strict threshold. Our thresholding results were congruent with those from an independent reporter of GABAergic neurons (*gad1b:GAL4, UAS:GCaMP6s*), supporting the validity of the immunostaining results (Fig. 3E). We found that only 0.4% of neurons were GABAergic and 1.4% of neurons were glycinergic (Fig. 3F, G). This meant that our cultures produced networks with an excitatory-to-inhibitory (E:I) neuron ratio of approximately 50:1, with a majority of inhibitory neurons being glycinergic. Previous studies have employed *in situ* hybridization and antibody labelling to support the idea that these ratios roughly reflect the neurotransmitter subtype composition of embryonic zebrafish brains (Macdonald et al., 1994, Martin et al., 1998, Cui et al., 2005, Higashijima et al., 2004). Thus, it appears likely that the sparseness of inhibitory neurons in our cultures is a product of their sparseness *in vivo*. Although this does not reflect the eventual neurotransmitter type composition in larvae and adults (Brenet et al., 2019, Pedroni and Ampatzis, 2019), it does provide confidence that our cultures can be used to study developmental and fundamental neuronal processes relevant to the *in vivo* embryonic context.

To test whether neurons in our pure cultures formed synaptic connections, we employed antibodies against SV2, PSD-95, and Gphn, which label presynaptic vesicles, excitatory postsynaptic densities, and inhibitory postsynaptic densities, respectively. By setting strict criteria for what constituted a synapse within a 3D confocal stack, we confirmed that our protocol produced synaptically interconnected neuronal networks with both excitatory and inhibitory connections. We found that there were, on average, more excitatory connections than inhibitory ones, which is consistent with our finding that a minority of neurons populating the cultures were inhibitory. Interestingly, there appeared to be a discrepancy between the synaptic E:I ratio and the neuronal E:I ratio. The synaptic E:I ratio averaged just under 2.5:1, whereas the neuronal E:I ratio was approximately 50:1. This finding implies that inhibitory neurons, on average, formed roughly twenty times more synaptic connections than excitatory neurons. This extensive connectivity of inhibitory neurons may represent underlying activity-dependent homeostatic drive to equilibrate network physiology and avoid hyperexcitability. Given that neurodevelopmental disorders are overrepresented by genes affecting inhibitory neurons (Ali Rodriguez et al., 2018), employing our culturing protocol may provide a robust platform upon which to explore developmental windows where inhibitory neurons are strongly outnumbered by excitatory neurons.

Lastly, we used our optimized protocol to perform phenotyping with cultures drawn from genetic mutants that model human diseases. We cultured neurons from embryos with presumed null mutations in the voltage-gated sodium channel subunit *scn1lab*. We chose *scn1lab* because its disruption is known to produce extensive neuropathology in zebrafish larvae (Banerji et al., 2021; Baraban et al., 2013; Brenet et al., 2019; Schoonheim et al., 2010; Tiraboschi et al., 2020). As such, it was likely to produce changes in our cultures, provided that the cultures recapitulated the relevant *in vivo* neurobiology. We found that cultures grown from *scn1lab*^−/−^ animals trended towards having fewer GABAergic neurons (Fig. 3F), which is consistent with known effects in *scn1lab* mutant zebrafish (Brenet et al., 2019; Tiraboschi et al., 2020). The number of glycinergic neurons did not appear to be affected by loss of *scn1lab* (Fig. 3G), indicating specificity towards GABAergic neurons. This finding is consistent with studies in rodents that highlight the importance of *SCN1A* expression in GABAergic neurons (Han et al., 2012, Ogiwara et al., 2007). Interestingly, there was no measurable change in the synaptic E:I ratio between cultures sourced from *scn1lab* animals and their WT siblings (Fig. 4G), indicating possible compensation by the remaining inhibitory neurons.

We also found a novel phenotype in cultures containing FBS, which were not purely populated by neurons. Cultures from *scn1lab*^−/−^ animals contained a surprisingly high number of non-neuronal cells (Fig. 2E, F). The presence of some of these cells, such as those morphologically resembling Schwann cells or oligodendrocytes, can only be explained by neural crest cells (NCCs), or progenitors of NCCs, proliferating and differentiating *in vitro*. As such, our findings suggest that *scn1lab* plays a role in suppressing proliferation of NCCs or cells from which NCCs are derived. Alternatively, *scn1lab* may promote differentiation of non-NCC lineages such that when it is lost, NCCs or their progenitors become more likely to proliferate and differentiate into NCC-related lineages. RNAseq of *scn1lab*^−/−^ brain tissue from 6 dpf larvae provided transcriptomic confirmation of the *in vitro* observations by revealing downregulation of the neuronal transcripts *slc17a6b* and *gad1b* and upregulation of the melanocyte markers *tyrp1a* and *tyrp1b* (Figure 5). The specific increase in melanocyte markers is particularly meaningful in the context of increased NCC-lineage cells in *scn1lab*^−/−^ cultures given that melanocytes are derived from NCCs (Dawes and Kelsh, 2021). One limitation of our transcriptomic analysis is that it was performed on tissue that included dermal and brain tissue. Because *scn1lab* mutants are hyperpigmented, it is possible that the signatures indicating increased abundance of melanocytes originate from the dermal tissue within our samples. Although the exact tissue origin of the melanocytes is an important consideration, the presence of these cells alone is sufficient to confirm the usefulness *in vitro* cultures with *in vivo* phenomena considering they are derived from NCCs. Our findings are also in agreement with other another study that applied RNA-seq to *scn1lab* mutant zebrafish larvae, which uncovered an enrichment of radial glia progenitors in *scn1lab* mutant brains (Tiraboschi et al., 2020). As such, our *in vitro* platform appropriately models phenomena that occur *in vivo* and, in so doing, provides an additional framework to explore the underlying mechanisms of autism, schizophrenia, and other psychiatric conditions.

## Acknowledgments

We thank the members of the Scott lab for guidance of this project and feedback on the manuscript. Support was provided by two Simons Foundation Research Awards (625793 to EKS and 573508 to EJH and EKS), two ARC Discovery Project Grants (DP220103812 and DP230102614) and an NHMRC Investigator Grant (2027072) to EKS. The research reported in this publication was supported by the National Institute of Neurological Disorders and Stroke of the National Institutes of Health under Award Number R01NS118406 to EKS. The content is solely the responsibility of the authors and does not necessarily represent the official views of the National Institutes of Health.

